# Convergent evolution of “genome guardian” functions in a parasite-specific *p53* homolog

**DOI:** 10.1101/2021.11.28.470261

**Authors:** George Wendt, Divya A. Shiroor, Carolyn E. Adler, James J. Collins

## Abstract

**Significance:** P53 is a widely studied tumor suppressor that is found throughout metazoans, including invertebrates that do not develop malignancies. The prevailing theory for why this is the case is that P53 originally evolved to protect the germline of early metazoans from genotoxic stress such as UV radiation. In this study, we examine the function of two P53 homologs in the parasitic flatworm *Schistosoma mansoni*. The first is orthologous to canonical P53, and regulates stem cell maintenance and differentiation. The second P53 gene is a parasite-specific paralog that is required for the normal response to genotoxic stress. This implies that the ability to respond to genotoxic stress in parasitic flatworms may have arisen from convergent evolution.

## Introduction

First described as a “guardian of the genome” in 1992(1), P53 has long been studied in the context of vertebrate cancer where its function and regulation is relatively well-understood(2). The P53 family of proteins, however, is widely conserved across metazoans, including invertebrates that do not seem to develop malignancies(3, 4). Studies in invertebrate model organisms have suggested that invertebrate P53 homologs generally respond to genotoxic stress by inducing cell death, much like vertebrate P53. In the absence of malignancies, this behavior is hypothesized to be instrumental in eliminating germ cells that have acquired mutations, supporting the idea that the ancestral function of P53 is to defend the integrity of the genome.

One problem with this inference of ancestral function, however, is that only a handful of invertebrate P53 homologs have been functionally studied to date. Though P53-like molecules have been shown to act as “genome guardians” (i.e., respond to genotoxic stress by inducing cell death) in invertebrates including *C. elegans*(5), *D. melanogaster*(6), and *N. vectensis*(7), examination of Platyhelminthes (flatworms) has been limited to the planarian flatworm *Schmidtea mediterranea* where the role of P53 in response to genotoxic stress was not thoroughly investigated(8, 9). The primary function of the planarian P53 homolog (SMED-P53) appears to be regulating the stem-cell mediated production of epidermal cells. RNAi-mediated loss-of-function of *Smed-p53* results in the loss of expression of the flatworm-specific transcription factor *zfp-1*(9), a known regulator of skin production in free-living and parasitic flatworms(10, 11) that is required for normal skin production in both worms.

Flatworm skin production is fascinating from both an evolutionary as well as a medical perspective. This is because of the clade Neodermata (literally “new skin”), the group of Platyhelminthes that contains virtually all parasitic flatworms, is united by the presence of a skin-like tissue known as the tegument. Unlike free-living flatworms that possess a simple multicellular epidermis, the tegument is a syncytial tissue that covers the entire surface of the organism, acting as an important interface between the parasite and the host(12). The tegument is known to play roles critical for survival inside the harsh environment of a host. In schistosomes, the tegument is involved in evading the hosts’ immune system, preventing hemostasis, and acquiring nutrients(13–16). Even more remarkable, throughout the course of evolution, tapeworms lost their gut in favor of absorbing nutrients through their tegument(17). That the tegument appears along with the adaptation to a parasitic lifestyle in flatworms suggest that the tissue may have been a key driving force that enabled these flatworms to become some of nature’s most prolific parasites(18). Furthermore, because the tegument plays critical roles in processes essential for parasite survival, better understanding how the tegument is made and maintained could lead to the development of new treatment and prevention methods to combat these deadly parasites, which are responsible for hundreds of thousands of deaths, billions of dollars in damage, and unquantifiable morbidity, primarily in the developing world(19).

Schistosomes are the most medically-important parasitic flatworm, killing over 200 thousand people every year and infecting over 200 million, with morbidity comparable to leishmaniasis, trypanosomiasis, and even malaria (20). Recent studies of the schistosome tegument and planarian epidermis demonstrated that despite having vastly different tissue organization, both animals have several commonalities in terms of skin production(9–11). Both planarians and schistosomes possess somatic stem cells termed neoblasts that constantly give rise to progenitor cells which migrate through the worm’s parenchyma until they eventually become part of the organisms’ epidermis or tegument, respectively. Aside from these anatomical similarities, both animals also rely on the flatworm specific transcription factor *zfp-1* in order to produce their skin. The presence of these similarities despite millions of years of evolutionary distance, unique tissue structures, and vastly different natural histories suggests that other factors that regulate epidermis production in planarians may be functionally conserved in schistosomes.

To that end, we decided to investigate homologs of *Smed-p53* in schistosomes. In doing so, we found two *P53* homologs in the schistosomes genome which we refer to as *p53-1* and *p53-2*. *p53-1* is orthologous to *Smed-p53* (as well as other ‘canonical’ P53 molecules such has human *TP53*) and appears to be functionally analogous to *Smed-p53*. *p53-2*, however, is a paralog of *Smed-p53* that appears to have arisen from a gene duplication event that occurred in the Neodermata and functions more akin to a “genome guardian”. Closer examination of *Smed-p53* suggests that the function of *p53-2* is novel and represents convergent evolution of genome guardian function in a P53 homolog, raising questions about the ancestral function of this important gene.

## Results

The *Schistosoma mansoni* genome contains two genes with the “p53 tumour suppressor family” InterPro ID (IPR002117)(21), Smp_139530 (henceforth *p53-1*) and Smp_136160 (henceforth *p53-2*). Both genes contain a p53 DNA binding domain (Pfam PF00870) but no apparent P53 transactivation domain (Pfam PF08563), P53 tetramerization domain (Pfam PF07710), or sterile alpha motif (Pfam 00536) (**Figure 1A**). Gene models from WormBase Parasite (Version WBPS16)(22, 23) identify 86 orthologs of *p53-1* including *H. sapiens TP53/TP63/TP73*, *D. melanogaster P53*, and *Smed-p53*, suggesting it represents the “canonical” P53 (**Figure 1B, Figures S1, S2, Table S1**). Examination of *p53-2*, however, identifies only 17 orthologs, all of which are only present in trematodes (flukes). Closer examination of other flatworms reveals that at least one apparent ortholog (identified via reciprocal BLAST) of *p53-2* is also present in 15 out of 16 cestode (tapeworm) genomes examined but not in any of the 2 monogenean or 19 free-living flatworm genomes examined (**Table S2**). Together, these data suggest a gene duplication event occurred in the common ancestor of Neodermata, resulting in *p53-2* representing a parasite-specific P53 paralog (**Figure 1C**). These data must be cautiously interpreted, however, due to the limited number and quality of flatworm genomes available. All *p53-1/p53-2* orthologs and genome/transcriptome databases referred to are listed in **Table S1** and **Table S3**, respectively.

**Figure 1.**
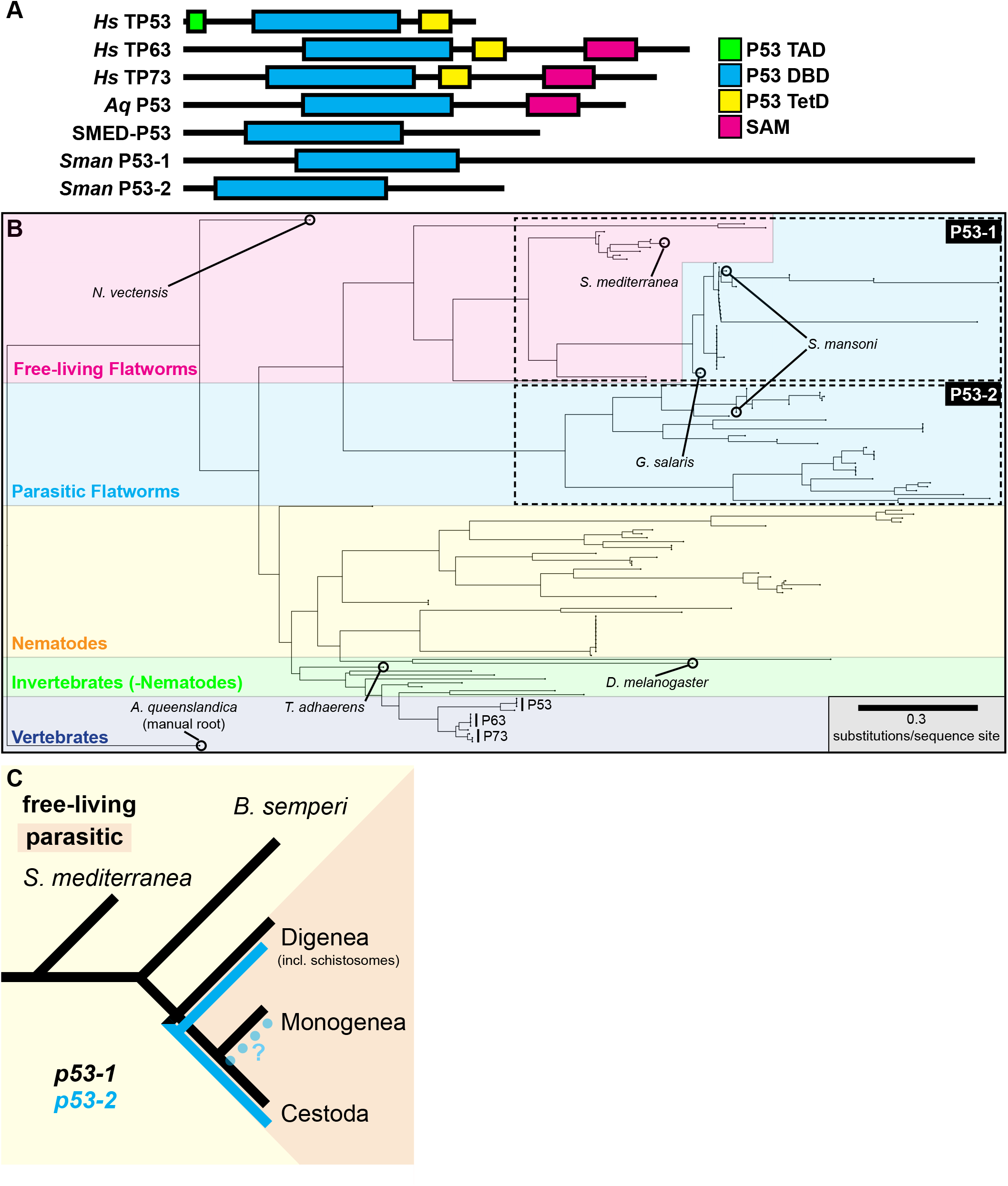
Phylogenetic analysis of p53-1 and p53-2 orthologs. (A) Schematic of domain structure of human TP53/TP63/TP73, *Amphimedon queenslandica* (*Aq*) P53, *S. mediterranea* SMED-P53, and S. mansoni p53-1/p53-2. TAD, transactivation domain; DBD, DNA binding domain; TetD, tetramerization domain; SAM, sterile alpha motif. (B) Phylogenetic trees of *p53-1* and *p53-2* orthologs, color-coded by remarkable clades. Flatworm p53-1 and p53-2 orthologs are indicated with dashed lines. (C) Cartoon depicting model of *p53-1* and *p53-2* evolution. Dotted line with question mark indicates the possible loss of *p53-2* orthologs in Monogenea.

Epidermal production in planarians mirrors tegument development in schistosomes in many ways, including how both processes are regulated by the same molecules(11). SMED-P53 is a known regulator of epidermal production in planarians(9, 10), so we hypothesized that one or both schistosome P53 homologs may also regulate tegument production. We began by examining the expression pattern of *p53-1* and *p53-2. SMED-P53* is expressed in planarian neoblasts and epidermal progenitors(8), so a functionally-conserved molecule would likely be expressed in analogous cells. Whole-mount *in situ* hybridization (WISH) of schistosome *p53* homologs (**Figure 2A**) revealed a punctate pattern of gene expression for *p53-1*, which is similar to other genes expressed in schistosome neoblasts and tegument progenitor cells(11, 24, 25). *p53-2*, however, had a more diffuse expression pattern in addition to apparent enrichment in the parasite’s gut and reproductive organs. Double fluorescence in situ hybridization (FISH) experiments demonstrated that *p53-1* is indeed expressed in schistosome neoblasts and tegument progenitor cells (**Figure 2B, Figure S3A**). *p53-2* is also expressed in many of these cells (**Figure 2B, Figure S3B**) in addition to in many other cell types including gut and reproductive cells (**Figure S3B**). We also examined the expression patterns of *p53-1* and *p53-2* using a recently published single-cell RNAseq atlas(25, 26) (**Figure 2C**) and found that the single-cell RNAseq data generally agree with our WISH/FISH results. Together, all of these data show that *p53-1* has an expression pattern very similar to *SMED-P53* whereas *p53-2* expression is comparatively unique.

**Figure 2.**
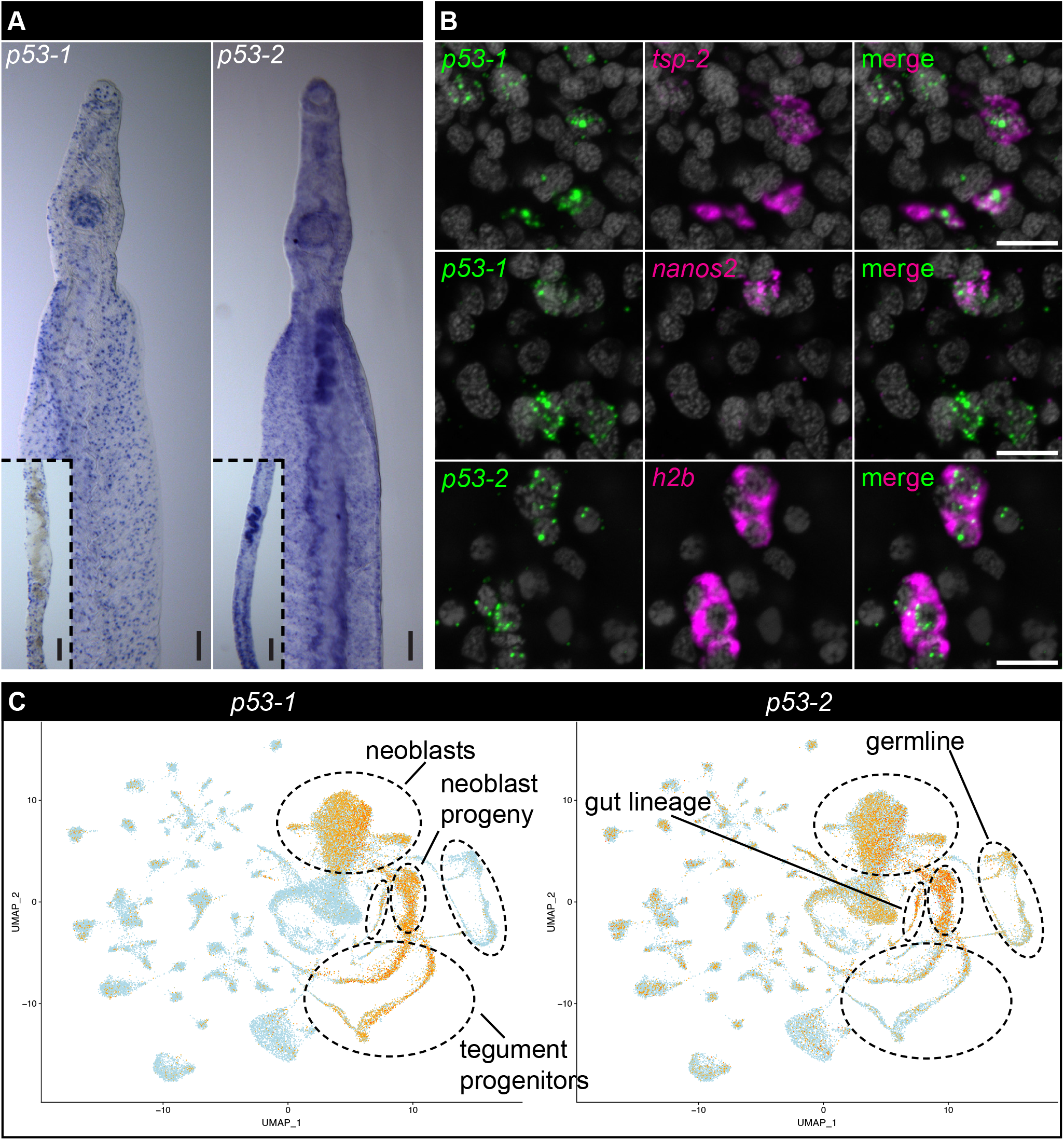
Expression pattern of schistosome p53 homologs. (A) Colorimetric WISH showing expression patterns of p53-1 and p53-2. (B) Double FISH experiment showing expression of *p53-1* relative to the tegument progenitor marker *tsp-2* and the neoblast marker *nanos2* as well as the expression of *p53-2* relative to the proliferative cell marker *h2b*. (C) Uniform manifold approximation plots showing expression patterns of p53-1 and p53-2 in adult schistosomes. Red indicates high expression, orange indicates medium expression, and blue indicates low/no expression. Important cell populations are indicated. Scale bars: (A) 100μm, (B) 10μm.

Next, we tested whether either schistosome P53 homolog was functionally related to SMED-P53 (i.e., a regulator of the “skin” lineage). *SMED-P53* RNAi eliminates epidermis-producing neoblasts leading to a loss of epidermal progenitors, ultimately resulting in impaired of epidermis production(9). As such, we examined the expression of *tsp-2* (a tegument progenitor marker, analogous to epidermal progenitors) and *eled* (a gut-producing neoblast marker) as well as EdU (a thymidine analog which labels proliferative cells) following RNAi of each P53 homolog. We found that *p53-1* RNAi results in complete loss of *tsp-2* cells and all *eled*-negative neoblasts, but no apparent change in the number of *eled*-positive neoblasts (**Figure 3A, Figure S4**). Loss of tegument progenitor cells and retention of gut-producing neoblasts should result in a loss of tegument production with no change in gut production, so we next examined tegument and gut production using EdU pulse-chase experiments. Consistent with loss of *tsp-2*-positive tegument progenitor cells but preservation of *eled*-positive gut producing neoblasts, *p53-1* RNAi resulted in complete loss of tegument production (**Figure 3B**), but did not cause any observable changes in gut production (**Figure 3C**). *p53-2* RNAi did not appear to have any effect on either gut or tegument production (**Figure 3, Figure S4**). Together, these data support a model where *p53-1* is a functional homolog of *SMED-P53* whereas *p53-2* appears to have a different role.

**Figure 3.**
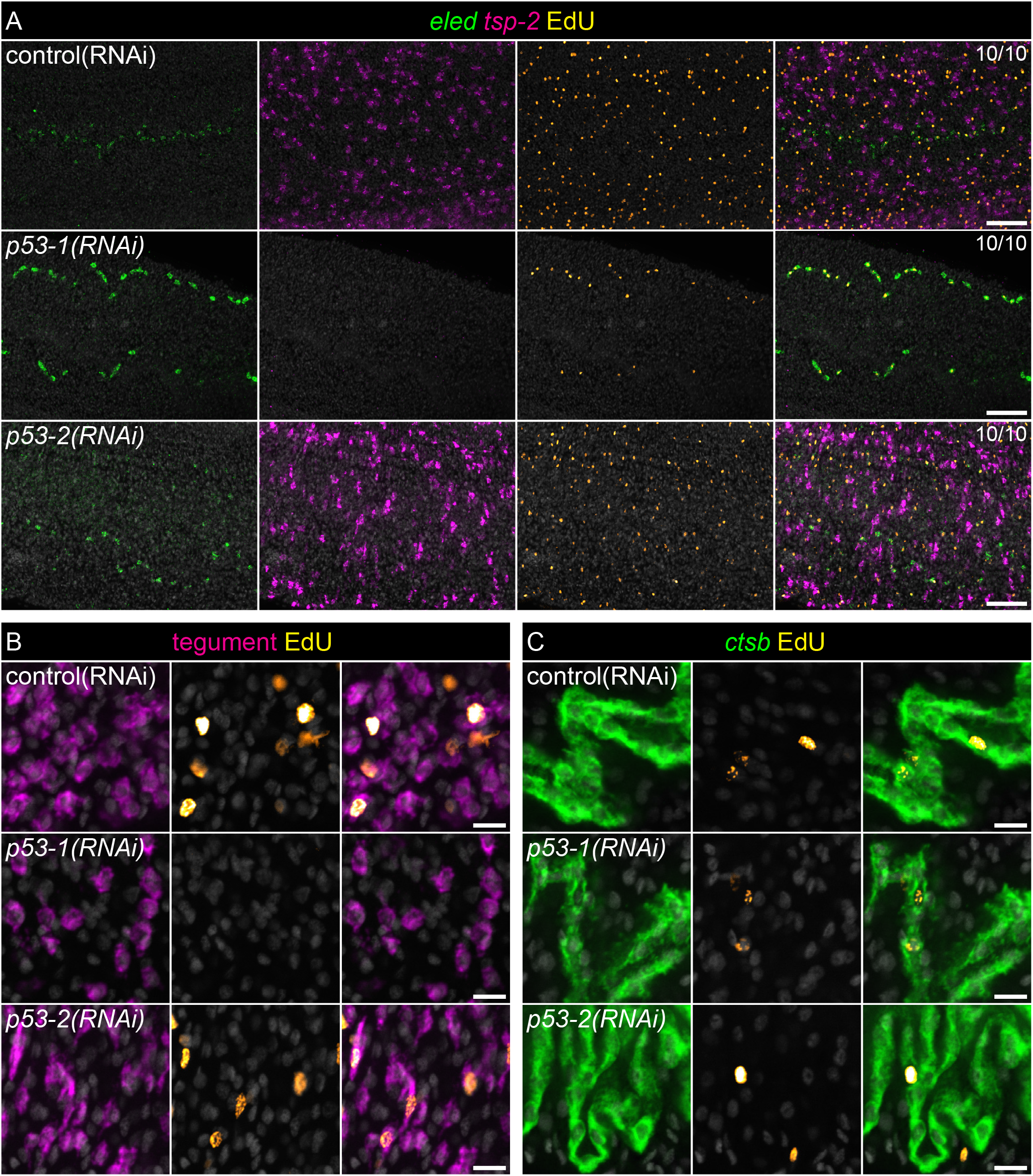
*p53-1* RNAi perturbs neoblast maintenance and differentiation. (A) Double FISH of the gut neoblast marker *eled* and the tegument progenitor marker *tsp-2* during *p53-1* and *p53-2* RNAi. Neoblasts are labeled with the thymidine analog EdU. (B) FISH of a cocktail of genes that mark the tegument during *p53-1* and *p53-2* RNAi. Neoblast progeny are labeled with EdU via a pulse-chase experiment. (C) FISH of the gut marker *ctsb* after *p53-1* and *p53-2* RNAi. Neoblast progeny are labeled with the EdU via a pulse-chase experiment. Fraction indicates number of worms that are similar to representative image. Scale bars: 50μm.

The prevailing theory as to the function of invertebrate P53 homologs is that they respond to DNA damage by eliminating affected cells(4). Given that *p53-2* had no apparent role in tegument production but is still a P53 homolog, we next tested whether *p53-2* has any role in the parasite’s response to genotoxic stress. A low dose (20gy) of radiation is sufficient to deplete the vast majority of proliferative cells in adult parasites (**Figure 4A**). *p53-1* RNAi offers no protection from this effect, but *p53-2* RNAi limited the impact of radiation on EdU^+^ proliferative cells (**Figure 4A**).

**Figure 4.**
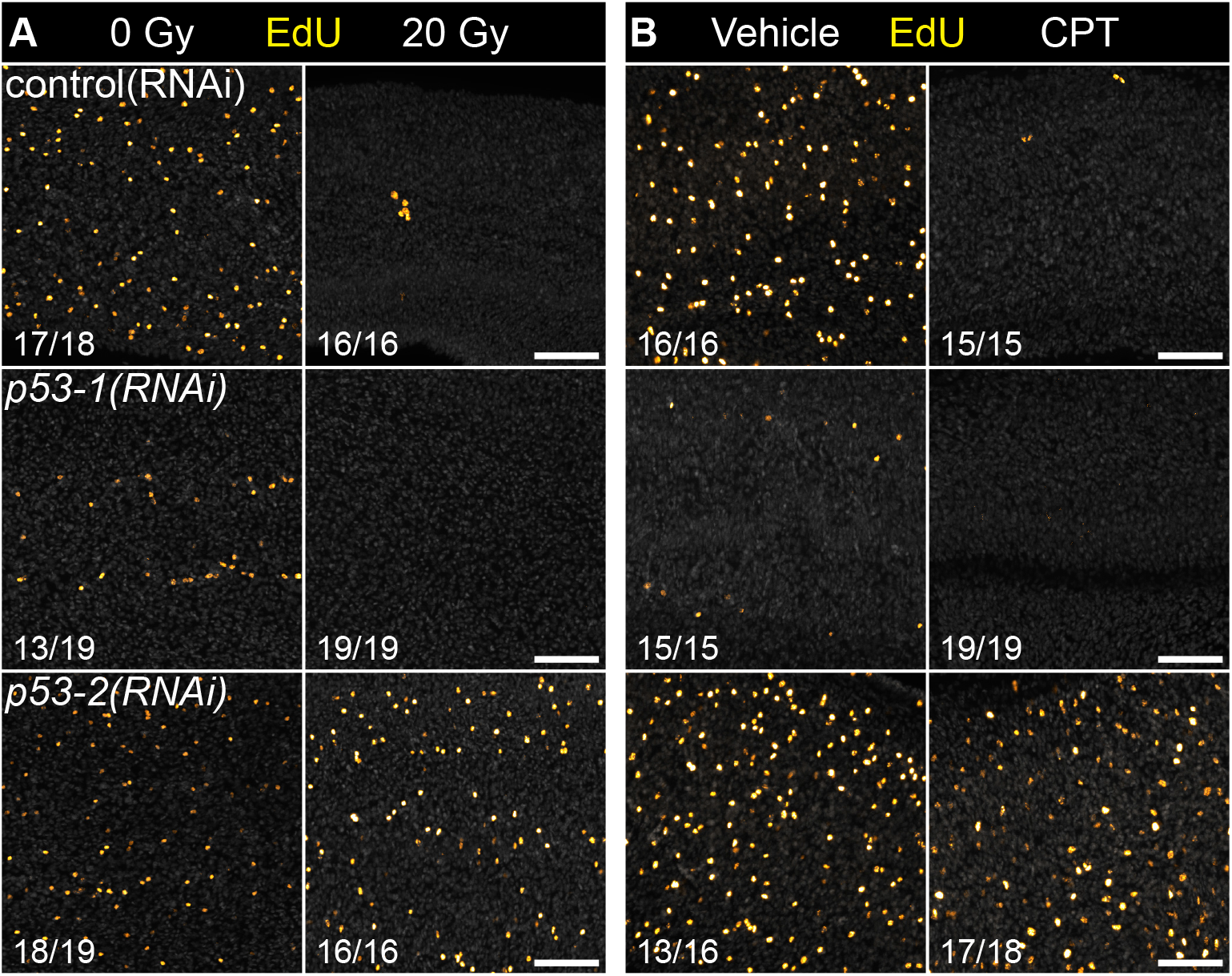
*p53-2* RNAi abrogates normal response to genotoxic stress. (A-B) EdU labelled neoblasts following irradiation (A) or cisplatin treatment (B) after *p53-1* or *p53-2* RNAi. (B). Fraction indicates number of worms that are similar to representative image. Scale bars: 50μm.

This protection from radiation extended beyond simply preserving EdU-positive proliferative cells. The EdU-positive neoblasts remaining after radiation still expressed markers of specialization (*eled*) and gave rise to tegument progenitors (**Figure S5**), suggesting that they are at least partially functional. *p53-2* RNAi also protected the parasite’s neoblasts from the effects of chemical genotoxic stress in the form of the DNA crosslinking agent cisplatin (CPT)(27) (**Figure 4B**), though CPT appeared to have a greater negative impact on production of *tsp-2*-positive tegument progenitor cells compared to radiation (**Figure S6**). These data support the existence of a genotoxic stress response function for *p53-2* that is not present in *p53-1*.

The ancestral function of P53 in regulating the response to genotoxic stress may have been lost or diverged between planarians and schistosomes. In planarians, radiation exposure rapidly induces body-wide apoptosis(28) and loss of Smed-p53 expression(8, 9), but the precise role of p53 in initiating apoptosis remains unresolved. To directly test whether SMED-P53 is required for stem cell death in response to DNA damage, we exposed *Smed-p53(RNAi)* animals to radiation and analyzed rates of apoptosis. Staining purified neoblasts with the apoptosis marker Annexin-V allows quantification of apoptosis within stem cells(29). Unirradiated *Smed-p53(RNAi)* animals showed no change in overall numbers of neoblasts or rates of apoptosis (**Figure 5**). Radiation exposure caused an equivalent depletion of neoblasts and corresponding increase in apoptosis as in control animals, indicating that *Smed-p53* is not required for apoptosis in response to genotoxic stress in planarians. Together, these results suggest the genotoxic stress response function of *p53-2* is in fact novel and the function of the *p53-1* ortholog in both schistosomes and planarians is to regulate neoblast survival and differentiation.

**Figure 5.**
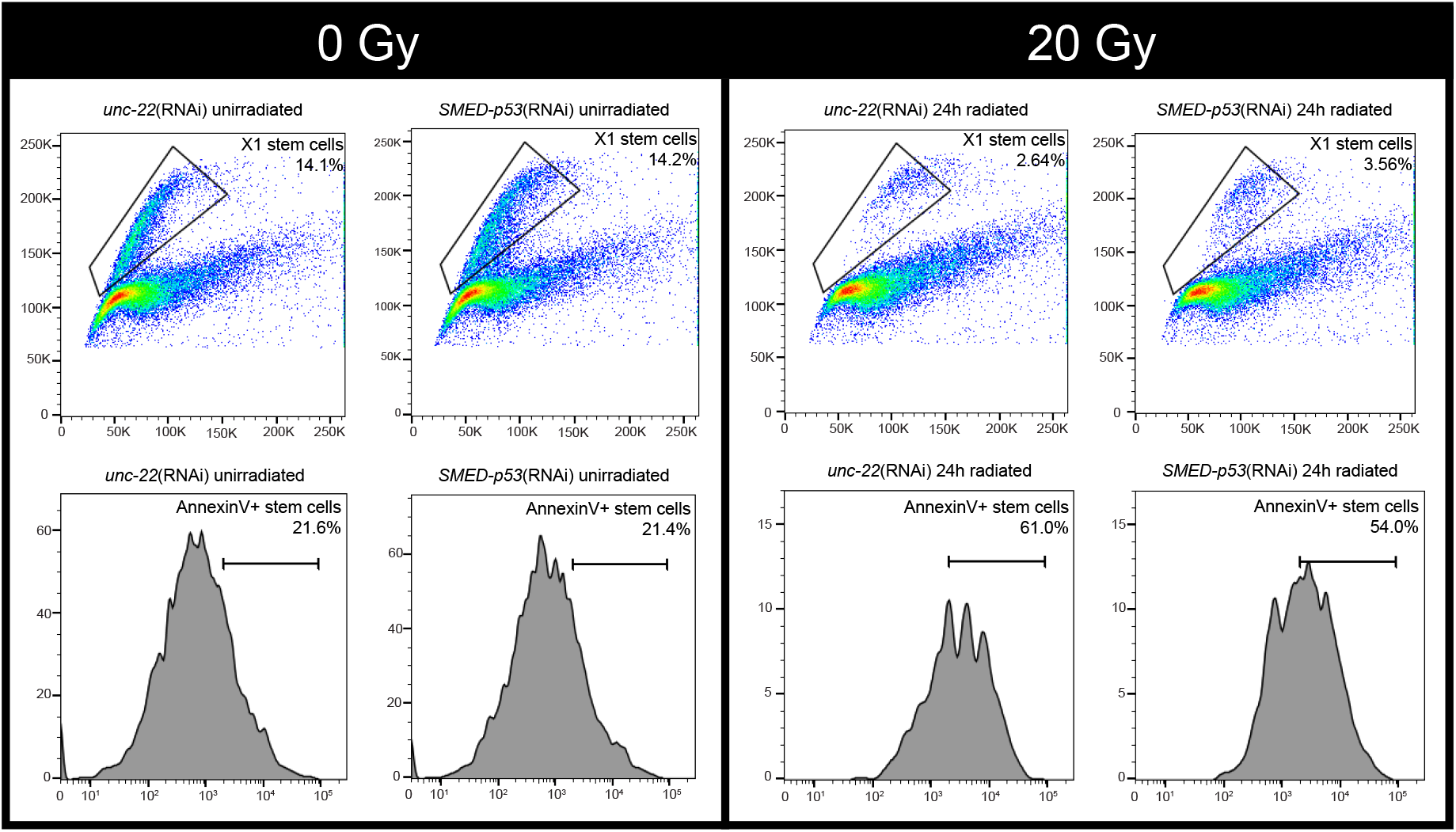
SMED-P53 RNAi does not protect planarian neoblasts from genotoxic stress. FACS plots of Annexin V flow cytometry experiment. Percentage of neoblasts remaining following irradiation (“X1 stem cells”) and percentage of apoptotic neoblasts (“AnnexinV+ stem cells”) is indicated in the 2D flow plot and flow histogram, respectively.

## Discussion

The evolution of the P53 family of proteins is both complex and fascinating. Based on data from the nematode *C. elegans*, insect *D. melanogaster*, and sea anemone *N. vectensis*, the ancestral role of P53 was likely to defend the germline from genotoxic stress(5–7). This stress could be in the form of exogenous radiation or endogenous transposable elements(30). This is an attractive hypothesis because it explains why invertebrates that do not appear to be susceptible to malignancies would possess a “tumor suppressor”(4).

Our studies in flatworms, however, suggest that the ancestral flatworm P53 (i.e. flatworm *p53-1* orthologs) functions in stem cell maintenance and differentiation as opposed to regulating the response to genotoxic stress. The expression pattern and RNAi phenotype of *p53-1* and *SMED-P53* are virtually identical, suggesting that *p53-1* orthologs may be functionally conserved across Platyhelminthes. Future investigation of *p53-1* orthologs (as well as *zfp-1* homologs) in other model flatworms will help clarify the extent of the evolutionary conservation of the regulation “skin” production and neoblast differentiation. *p53-2*, on the other hand, appears to be a more recent invention, emerging concurrently with Neodermata. Though further studies of *p53-2* orthologs in other flatworms is required, it is very tempting to speculate that the genotoxic stress response function in *p53-2* represents convergent evolution of the “genome guardian” function thought to be the original function of P53-family proteins. This raises two major questions: is the ancestral function of P53 actually stem cell regulation rather than the genotoxic stress response, or did that spontaneously appear in an early metazoan ancestor and simply persist because of its usefulness? Additionally, if *p53-2* is in fact a parasite-specific homolog that responds to genotoxic stress, why do parasites but not free-living flatworms need such a gene?

Ultimately, it is not possible to definitively answer fundamental questions regarding evolutionary history but we come closer to understanding things such as ancestral function with every new model organism that we functionally study. Given that our understanding of the ancestral function of P53 is based on only a handful of model organisms, it is perhaps not surprising that these data challenge the existing notion that the ancestral function is to respond to genotoxic stress. We should note that these data do not refute previous theories of ancestral function; several possible explanations for our data exist. The most obvious would be that we failed to identify any *p53-2* orthologs in free-living flatworms because we have not examined enough high-quality genomes. We surveyed 19 free-living flatworm genomes and failed to identify any *p53-2* orthologs, as opposed to parasitic flatworms where searching 35 genomes identified at least one *p53-2* orthologs in 31 of them. It is also possible that *S. mediterranea* is the outlier whose *p53-1* ortholog does not possess the genome guardian function. Together, this uncertainty underscores the importance of isolating and sequencing genomes of diverse species for the purpose of advancing our understanding of evolutionary and developmental biology. Another potential explanation for the appearance of a genotoxic stress-response function is that said function was lost early on in flatworm evolution but spontaneously reappeared in Neodermata. While this is not the most parsimonious explanation of our data, it is possible that P53-family proteins simply have a propensity for acquiring the ability to respond to genotoxic stress due to the nature of the DNA sequences that they bind. Indeed, if the ancestral function of P53 was in fact the regulation of stem cell maintenance and differentiation, then the ability to respond to genotoxic stress independently evolved at least twice (once in a basal ancestral metazoan and once again in parasitic flatworms).

The second question that arises from these studies is simpler but much harder to answer: why do parasites need a P53 homolog that responds to genotoxic stress? Unlike free-living flatworms, parasitic flatworms actually spend much of their life living inside of a host where they are presumably protected from environmental sources of genotoxic stress such as UV radiation. One factor to consider is the complex lifecycles that are characteristic of parasitic flatworms. Most alternate between at least one vertebrate and one invertebrate host, meaning that they are not always protected from environmental genotoxic stress (i.e. UV radiation). As such, *p53-2* might respond to experimental genotoxic stress in adult schistosomes, but physiologically it may only be important in other developmental stages that were not examined in this study. Another possible explanation pertains to endogenous threats to genome stability: transposable elements. *p53-2* orthologs could be involved in the suppression of transposable elements as has been demonstrated for P53 proteins in flies, zebrafish, and humans(30, 31). This hypothesis is especially attractive because many parasitic flatworm genomes studied to date appear to have lost components of the transposon-suppressing piRNA pathway(32, 33), raising questions as to how they protect their genome from transposable elements. Further investigation of transposon activity following *p53-2* RNAi may yield interesting results.

As more flatworms become available for study in the laboratory, we will be able to better answer these questions specifically, and possibly broader evolutionary questions as well. Despite being arguably the most widely studied gene in history, there is still a great deal that we do not know about how P53 came to be. Further studies of P53 homologs in flatworms could clarify the origins of this important gene while also teaching us more about the basic biology of parasitic flatworms, which will help in the development of novel therapeutics against these deadly pathogens.

**Figure S1.**
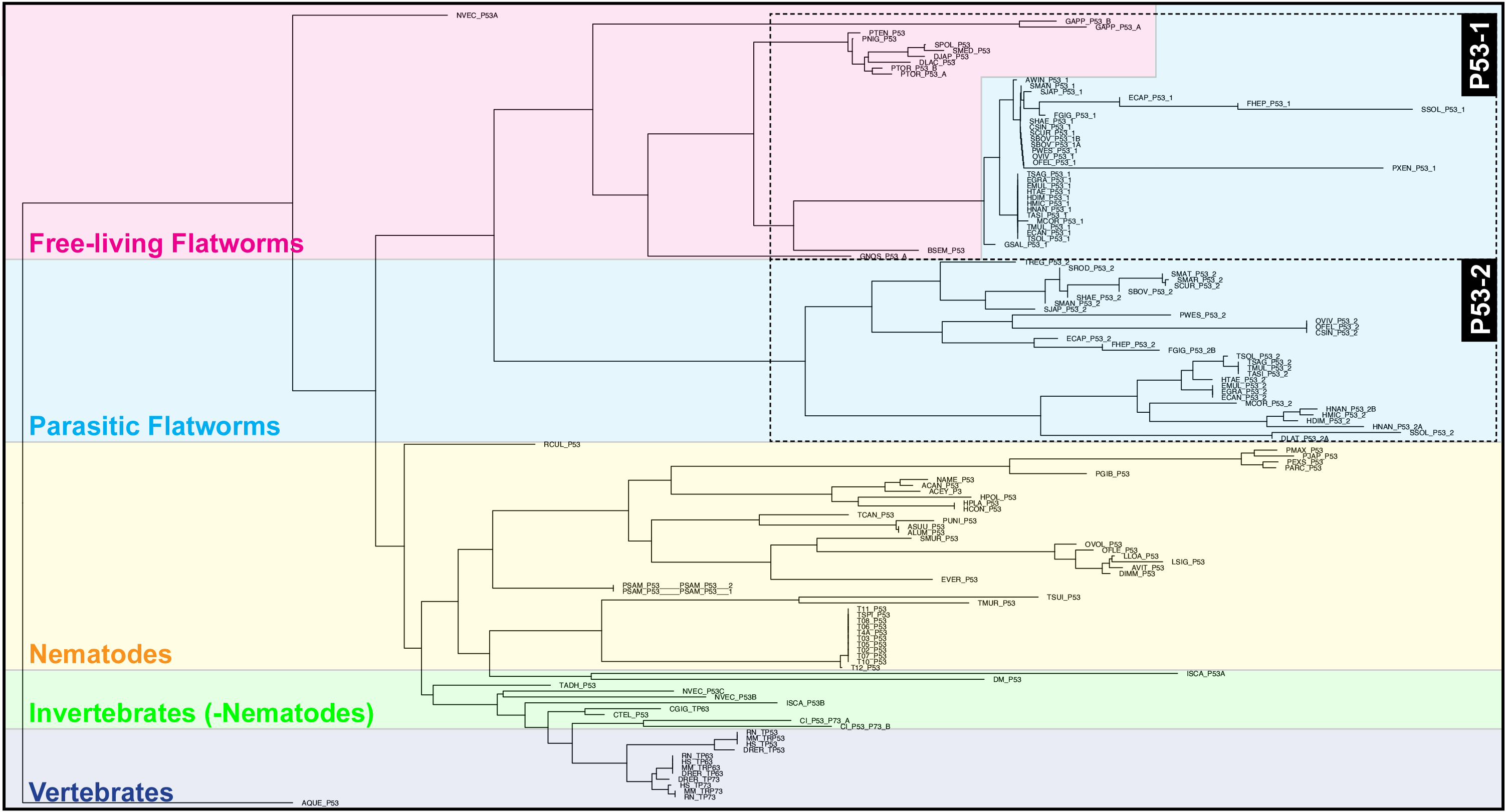
Phylogenetic tree of p53 homologs with tip labels. Phylogenetic trees of *p53-1* and *p53-2* orthologs, color-coded by remarkable clades with tip labels (species and homolog) included. Flatworm p53-1 and p53-2 orthologs are indicated with dashed lines.

**Figure S2.**
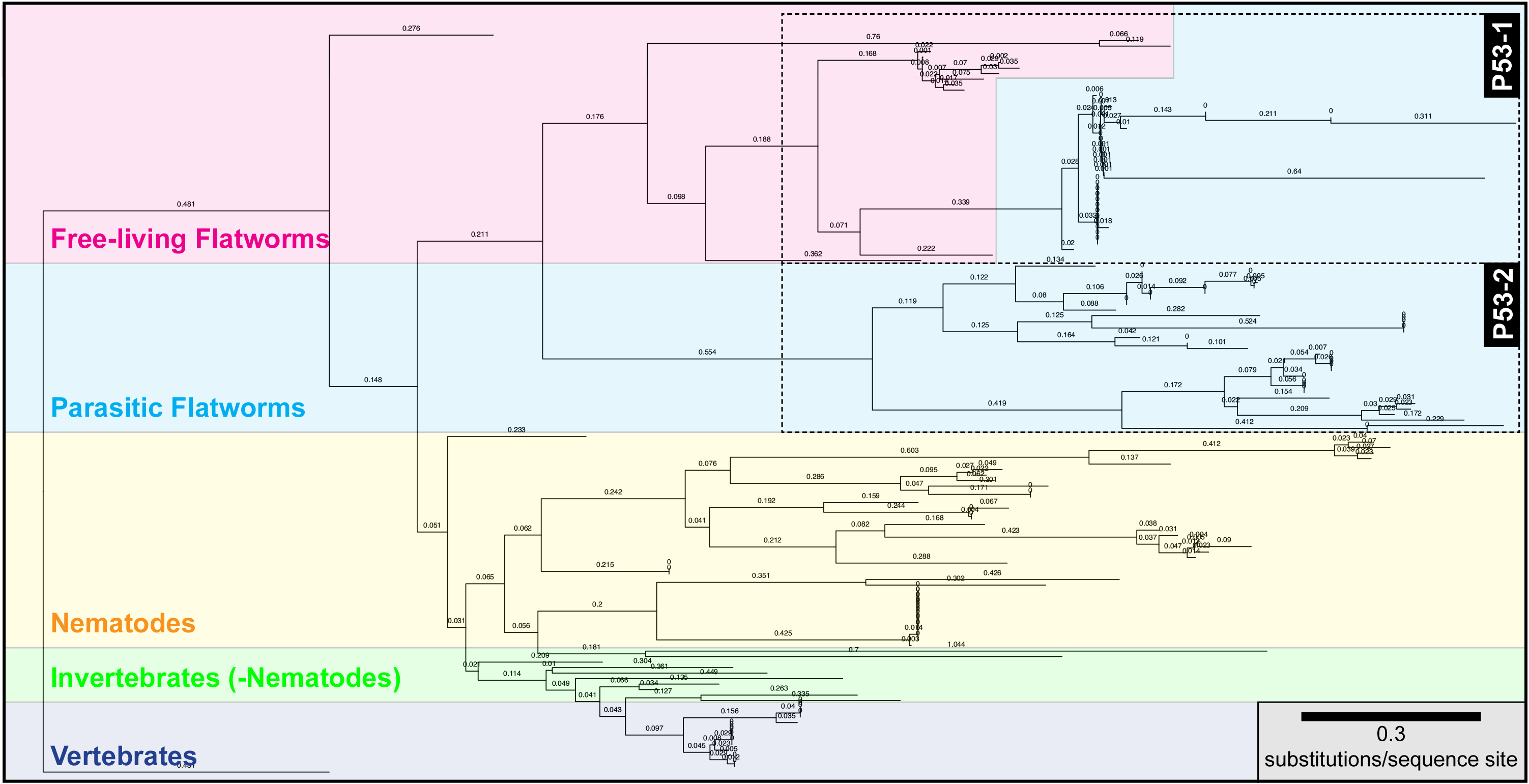
Phylogenetic tree of p53 homologs with branch lengths. Phylogenetic trees of *p53-1* and *p53-2* orthologs, color-coded by remarkable clades with branch lengths (substitutions per sequence site) included. Flatworm p53-1 and p53-2 orthologs are indicated with dashed lines.

**Figure S3.**
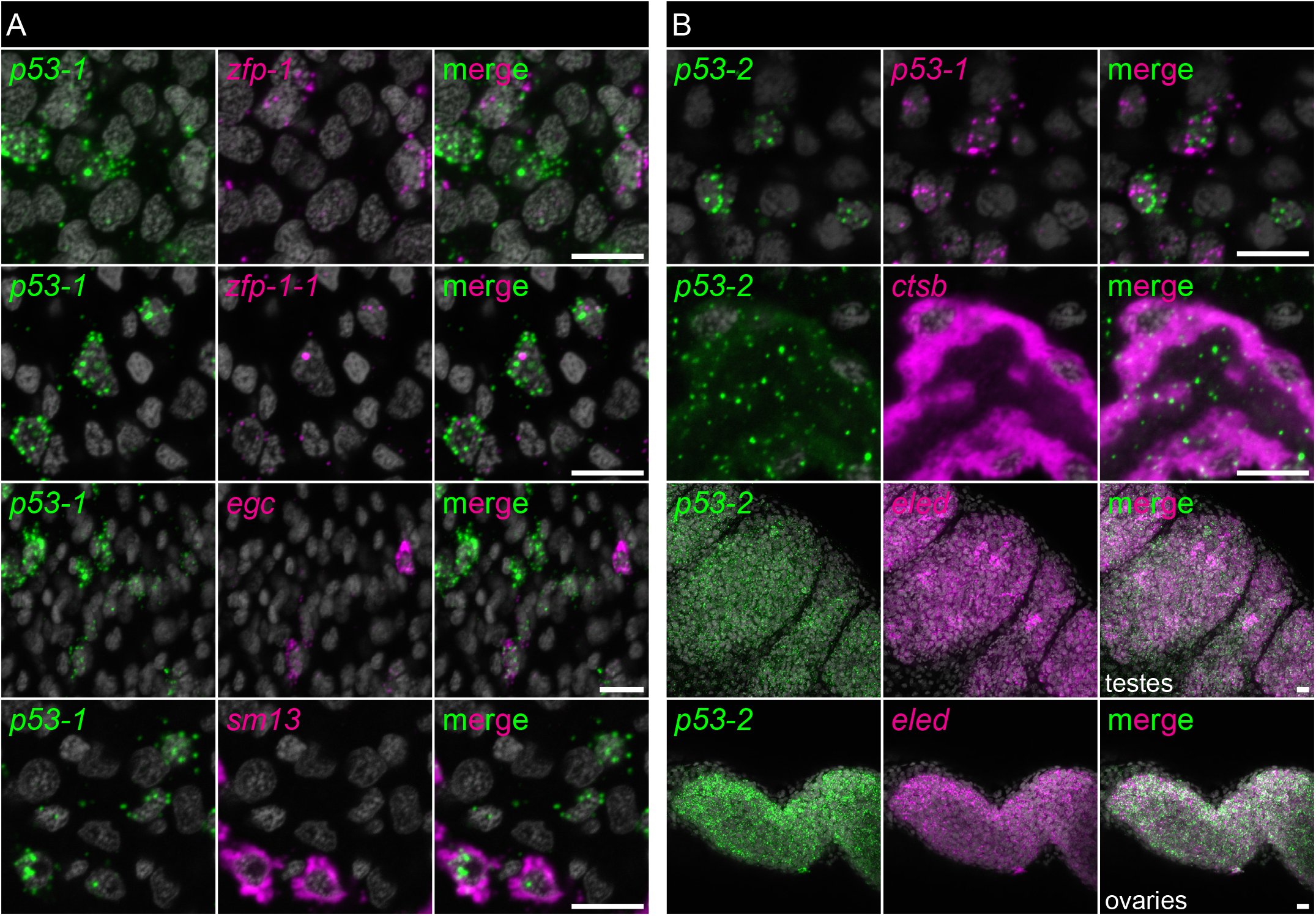
Expression pattern of schistosome p53 homologs. (A) Expression pattern of *p53-1* relative to the neoblast marker *zfp-1* and the tegument progenitor subset markers *zfp-1-1*, *egc*, and *sm13*. (B) Expression pattern of *p53-2* relative to *p53-1*, the gut marker *ctsb*, and the germ stem cell marker *nanos1* in the testes and the ovaries. Scale bars: 10μm.

**Figure S4.**
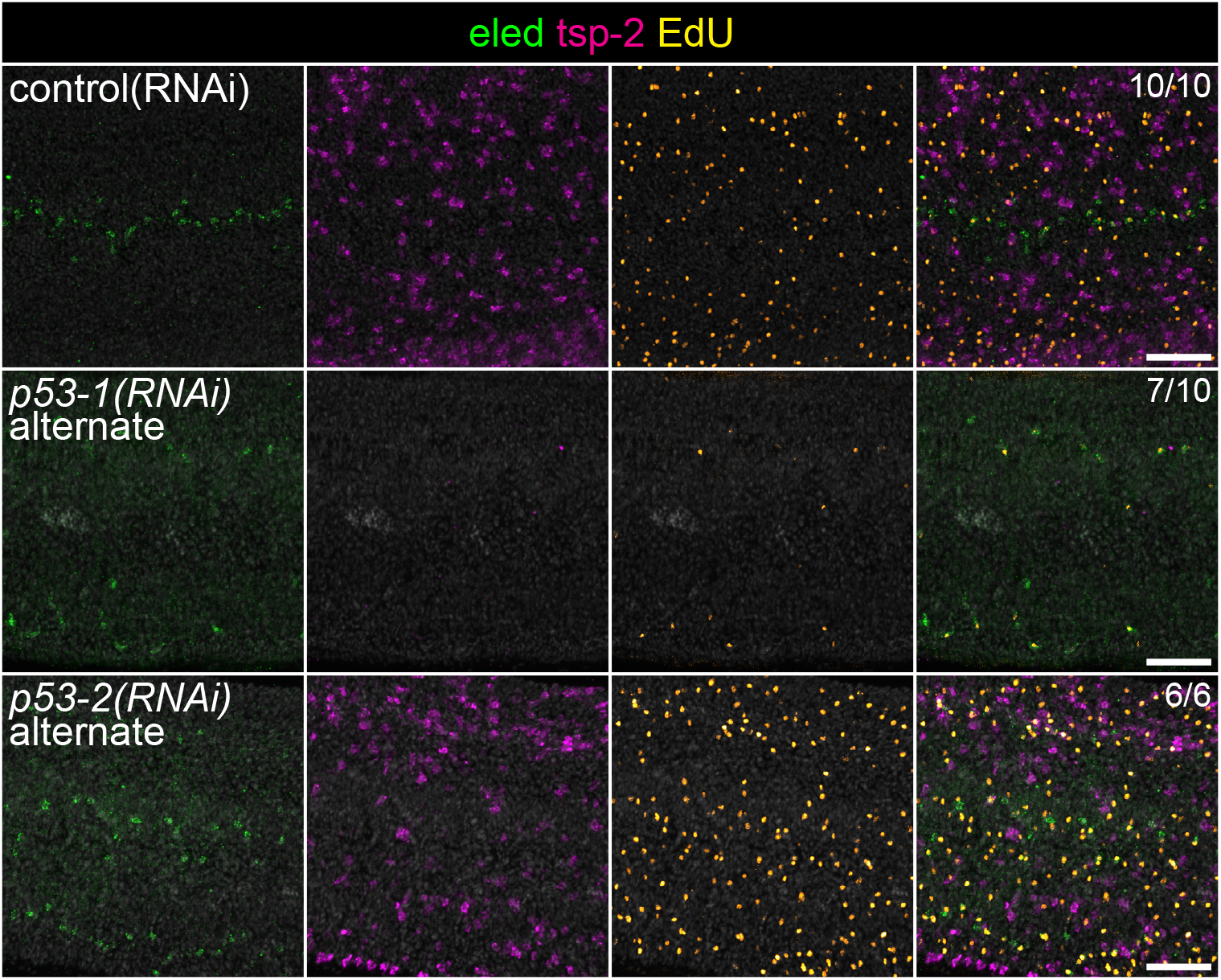
Non-overlapping RNAi constructs cause the same phenotype. Double FISH of the gut neoblast marker *eled* and the tegument progenitor marker *tsp-2* during *p53-1* and *p53-2* RNAi. Neoblasts are labeled with the thymidine analog EdU. Note that images for “control(RNAi)” are reproduced from Figure 3A to allow reference to control phenotype. All images shown are from the same biological replicate. Fraction indicates number of worms that are similar to representative image. Scale bars: 50μm.

**Figure S5.**
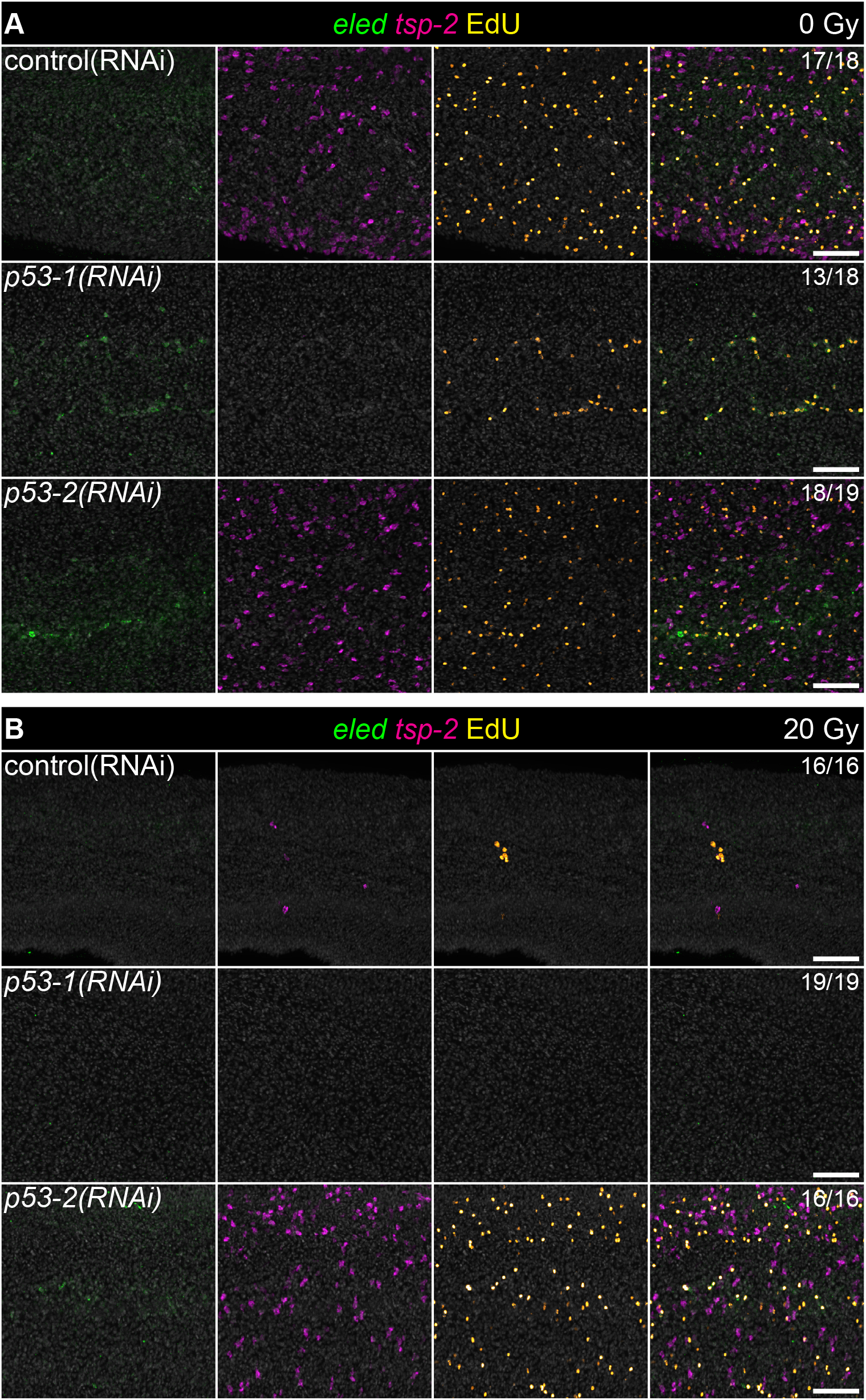
Neoblasts that persist after radiation in *p53-2* RNAi differentiate normally. (A-B) Double FISH of the gut neoblast marker *eled* and the tegument progenitor marker *tsp-2* after *p53-1* and *p53-2* RNAi with 0gy (A) or 20gy (B) of radiation. Neoblasts are labeled with the thymidine analog EdU. Fraction indicates number of worms that are similar to representative image. Scale bars: 50μm.

**Figure S6.**
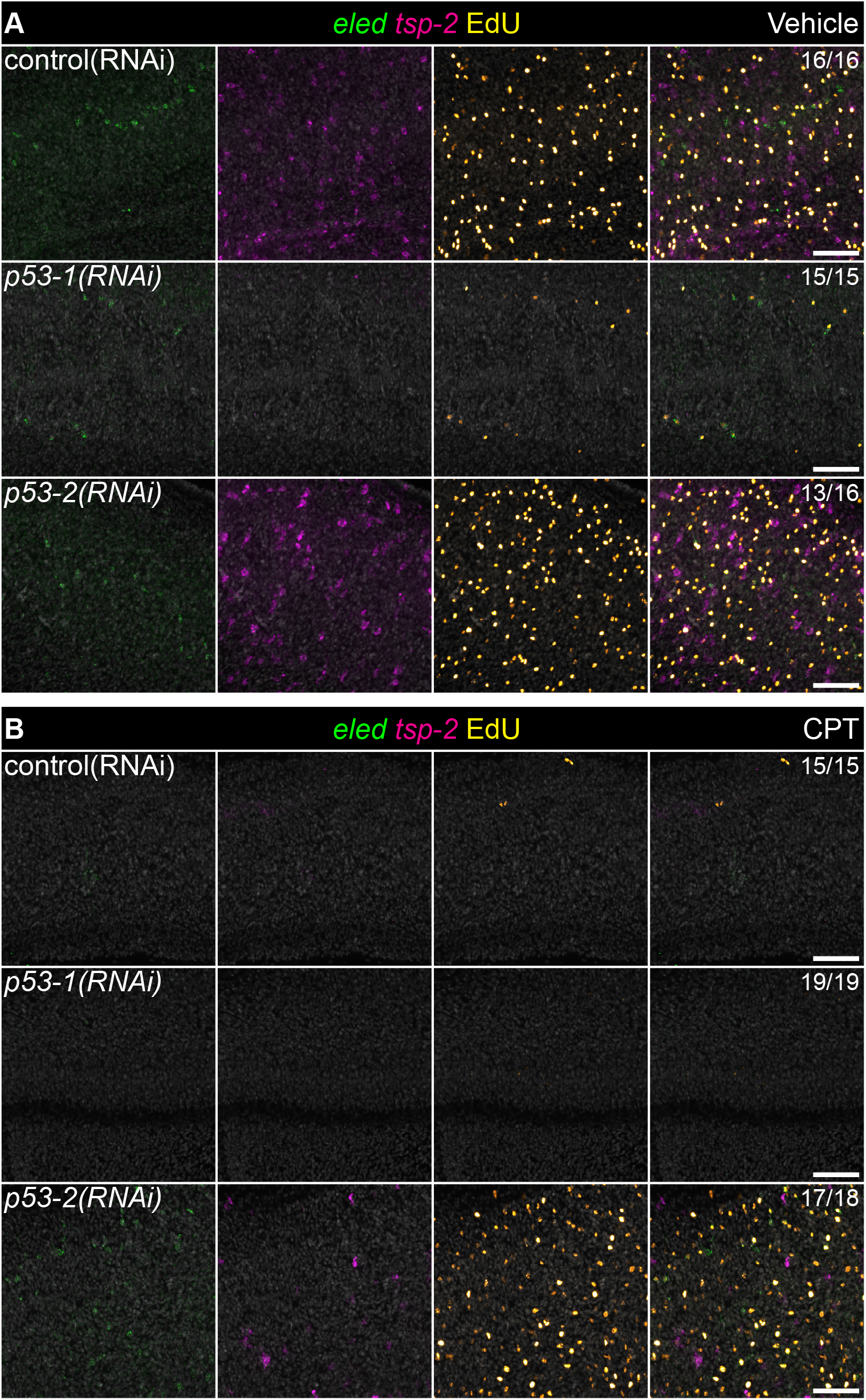
Neoblasts that persist after cisplatin treatment in p53-2 RNAi have differentiation defects. (A-B) Double FISH of the gut neoblast marker *eled* and the tegument progenitor marker *tsp-2* after *p53-1* and *p53-2* RNAi with vehicle (A) cisplatin treatment (B). Neoblasts are labeled with the thymidine analog EdU. Fraction indicates number of worms that are similar to representative image. Scale bars: 50μm.

## Materials and Methods

### Phylogenetic analysis

Identification of orthologs of schistosome P53 homologs was carried out using Wormbase Parasite (version WBPS16)(22, 23). Examination of *p53-2* (Smp_136160) orthologs initially only revealed trematode orthologs of *p53-2*. A reciprocal BLAST search of the *Hymenolepis diminuta* genome identified the *p53-2* ortholog WMSIL1_LOCUS11874. Wormbase Parasite then identified 15 orthologs of WMSIL1_LOCUS11874 in 14 unique cestode genomes, each of which was confirmed to be a *p53-2* ortholog via a reciprocal BLAST search. PlanMine v3.0 (https://planmine.mpibpc.mpg.de/planmine/begin.do) was additionally utilized in order to search 19 additional free-living flatworm genomes for *p53-2* orthologs. Domain structure of P53 homologs was obtained using SMART v9.0 (http://smart.embl-heidelberg.de/)(34, 35). Phylogenetic trees were generated using the FastME/OneClick Workflow function with default settings at NGPhylogeny.fr(36).

All *p53-1/p53-2* orthologs are listed in **Table S1**, organisms examined and the genome/transcriptome databases referred to are listed in **Table S3**, and the submission to NGPhylogeny is provided in **Table S4**.

### Parasite acquisition and culture

Adult S. mansoni (NMRI strain, 6–7 weeks post-infection) were obtained from infected female mice by hepatic portal vein perfusion with 37°C DMEM (Sigma-Aldrich, St. Louis, MO) plus 10% Serum (either Fetal Calf Serum or Horse Serum) and heparin. Parasites were cultured as previously described(24). Unless otherwise noted, all experiments were performed with male parasites. Experiments with and care of vertebrate animals were performed in accordance with protocols approved by the Institutional Animal Care and Use Committee (IACUC) of UT Southwestern Medical Center (approval APN: 2017-102092).

### Planarian husbandry

Asexual *Schmidtea mediterranea* (strain CIW-4) stocks were maintained in 1x Montjuic water and fed homogenized beef liver as previously described (37). Worms were kept at 20°C in a shaded environment with a 12 hour light-dark cycle.

### Radiation

#### Schistosome radiation

Schistosomes were exposed to a 2000 rads dose of radiation using a Precision X-Ray (Connecticut, USA) X-Rad320 system. All animals were rinsed and placed in fresh Basch media following radiation.

#### Planarian radiation

Planarians were exposed to a 2000 rads dose of radiation by using a J.L. Shepherd & Associates (California, USA) Mark I-68 Irradiator. All animals were rinsed immediately after radiation and planarian water was replaced.

### Fluorescence Activated Cell Sorting

Annexin V Flow cytometry of Hoechst-stained cells was conducted as previously described (29, 38, 39) with some modifications. 30 animals per group were dissected in CMFB (CMF containing 0.5% BSA) and dissociated using 1:50 Liberase (2.5mg/mL, Roche 5401135001) in CMFB at 30°C with agitation at 300 rpm for 30 minutes on an Eppendorf ThermoMixer. Samples were triturated every 5 minutes with a pipette to aid dissociation. Dissociated cells were then diluted with equal volume of CMFB and pelleted by centrifugation (500 g, 5 minutes, RT). Pellets were resuspended in 1mL of CMFB and passed through a 30mm cell strainer (BD 340627). Strained cells were counted on an automated cell counter (Bio Rad TC20) and 2.83x 10^6^ cells/group were stained with 5μg/mL Hoechst 33342 (ThermoFisher H3570) in CMFB for 70 mins in the dark with gentle agitation. Cells were subsequently pelleted as before, and Hoechst solution was replaced with Annexin V staining buffer (2μL Annexin V-APC (Thermo Fisher A35110) diluted in 100μL freshly made 1X Annexin V buffer from a 10X stock solution (0.1M HEPES pH 7.4, 1.4M NaCl, and 25 mM CaCl_2_)). After staining for 15 minutes at RT, 400μL of 1X Annexin V buffer with 1μg/mL Propidium Iodide (Sigma P4170) was added to each tube. Cells were analyzed on a BD FACSymphony Analyzer. Data was analyzed in FlowJo (TreeStar, Ashland, OR).

### Labeling and imaging

Schistosome colorimetric and fluorescence *in situ* hybridization analyses were performed as previously described(11, 24, 40) with the following modification. To improve signal-to-noise for colorimetric in situ hybridization, all probes were used at 10 ng/mL in hybridization buffer. *in vitro* EdU labeling and detection was performed as previously described(40). All fluorescently labeled parasites were counterstained with DAPI (1 μg/ml), cleared in 80% glycerol, and mounted on slides with Vectashield (Vector Laboratories).

Confocal imaging of fluorescently labeled samples was performed on a Nikon A1 Laser Scanning Confocal Microscope. Unless otherwise mentioned, all fluorescence images represent maximum intensity projection plots. To perform cell counts, cells were manually counted in maximum intensity projection plots derived from confocal stacks. In order to normalize counts, we collected confocal stacks and normalized the number of cells counted to the length of the parasite in the imaged region. Brightfield images were acquired on a Zeiss AxioZoom V16 equipped with a transmitted light base and a Zeiss AxioCam 105 Color camera.

### RNA interference

#### Planarian RNAi

For SMED-P53 RNAi FACS experiments, RNAi was carried out as previously described(41). Briefly, double stranded RNA (dsRNA) was synthesized in vitro with PCR products for Smed-p53 and control gene unc-22 as a templates. Synthesized dsRNA was then mixed with a 4:1 liver:water paste containing 4μg of dsRNA per 10μL of liver. Animals were fed every 2 days, for a total of 3 feeds. Radiation was carried out 4 days after the last feed and animals were processed for Annexin V FACS 24h later.

#### Schistosome RNAi

All RNAi experiments utilized freshly perfused male parasites between 6-7 weeks post-infection (separated from females). dsRNA treatments were all carried out at 30 μg/ml in Basch Media 169. dsRNA was generated by *in vitro* transcription and was replaced daily for 3 days and then every 3 days until the end of the experiment. EdU pulses were performed at 5μM for 4 hours before either fixation or chase as previously described.

As a negative control for RNAi experiments, we used a non-specific dsRNA containing two bacterial genes(42). cDNAs used for RNAi and in situ hybridization analyses were cloned as previously described(42); oligonucleotide primer sequences are listed in **Table S5**.

### Statistical analysis

All two-way comparisons were analyzed using Welch’s t-test. The p-value for symbols denoting significance are noted in the figure legends.

